# Regulation of metal homeostasis by two F-group bZIP transcription factors bZIP48 and bZIP50 in rice

**DOI:** 10.1101/2023.05.18.541275

**Authors:** Tao Qing, Tian-Ci Xie, Qiao-Yun Zhu, Hai-Ping Lu, Jian-Xiang Liu

## Abstract

Zinc (Zn) is an essential microelement for plants as well as for human beings, and it regulates numerous metabolic process and protein activity. Zn deficiency not only impairs plant growth and development, but also affects human health. Rice (*Oryza sativa* L.) is a staple food for more than half of the world’s population, but how Zn homeostasis in rice is maintained is largely unknown. Here, we demonstrate that two F-group bZIP transcription factors, OsbZIP48 and OsbZIP50, are important for metal homeostasis in rice. Mutation of *OsbZIP48* and *OsbZIP50* impairs plant growth and reduces Zn, Fe, and Cu content in shoots. The N-terminus of either OsbZIP48 or OsbZIP50 contains two cysteine- and histidine-rich (CHR) domains, deletion or mutation of these CHR domains increases nucleus localization of OsbZIP48 and OsbZIP50. Both OsbZIP48 and OsbZIP50 have transcriptional activation activity, and the expression of 1117 genes involved in metal uptake, phenylpropanoid biosynthetic process, cell wall organization, et al., is reduced in *OsbZIP48* and *OsbZIP50* double mutant than that in wild-type ZH11 plant under Zn deficiency. Both OsbZIP48 and OsbZIP50 bind to the promoter region of the ZIP family transporter gene *OsZIP10* in rice, and activate the promoter activity of ZDRE *cis*-element derived from the *OsZIP10* promoter in effector-reporter assays. Mutation of the CHR domain of OsbZIP48 in *OsbZIP50* mutant background increases the content of Zn/Fe/Cu in brown rice seeds and leaves. Thus, this study reveals that OsbZIP48 and OsbZIP50 regulate metal homeostasis, especially under Zn deficiency in rice, and provides candidate target genes for biofortification of micronutrients in future.

**significance statement:** Zinc (Zn) is an essential microelement not only for plants but also for human beings. This paper shows that the N-terminal cysteine- and histidine-rich domain of OsbZIP48/50 is important for their nucleus localization, therefore transcriptional activity, and reveals the downstream genes of OsbZIP48/50 involved in metal homeostasis under Zn deficiency in rice.

## Introduction

Zinc (Zn) is one of the most important essential elements in plants, serving as a cofactor for numerous enzymes and regulatory proteins in plants (Broadley et al., 2012). Zn never undergoes redox state changes so that it doesn’t directly participate in electron transfer reactions, in contrast, it plays important roles in the detoxification of superoxide radicals, maintaining the membrane integrity, as well as in mediating enzymatic reaction, DNA-binding of transcription factors and protein–protein interactions (Clemens, 2022). Zn-deficit in soil is widely spread (Zeng et al., 2021), and it reduces crop yield and quality, as well as nutritional values, because Zn is also an essential element for human beings (Kawakami and Bhullar, 2018). Therefore, breeding crops with increased tolerance to zinc-deficit in soil should be on the agenda in the modern crop breeding programs.

Over the past few decades, several regulatory mechanisms underlying Zn homeostasis have been discovered in plants (Amini et al., 2022; Stanton et al., 2022). These include acquisition and uptake of Zn from soil, intracellular transport of Zinc across plasma membrane or organelle membranes, and translocation of Zn between different tissues/organs (Stanton et al., 2022). Among them, Zinc-Regulated Transporter, Iron-Regulated Transporter (ZRT-IRT)-like Protein (ZIP) family transporters are the highly conserved transporters for the uptake, intracellular transport, and detoxification of Zn^2+^, Fe^2+^, and Mn^2+^ in plants (Grotz et al., 1998). In rice (*Oryza Sativa*, L), there are 16 members in ZIP family, but only a few of them have been characterized to date. *OsZIP1* is abundantly expressed in roots and induced by excess Zn^2+^, Cu^2+^ and Cd^2+^ but not by Mn^2+^ and Fe^2+^, regulating metal efflux when Zn^2+^, Cu^2+^ and Cd^2+^ are excess in environment (Liu et al., 2019). In contrast, *OsZIP5* and *OsZIP9* are tandem duplicated genes which are induced in roots by Zn-limiting conditions (Lee et al., 2010a; Tan et al., 2020). OsZIP9 localizes at the root exodermis and endodermis, functioning as a major influx transporter of Zn^2+^ and other micronutrients (Huang et al., 2020; Tan et al., 2020; Yang et al., 2020). *OsZIP4* and *OsZIP8* are also induced by Zn deficiency in roots, analysis of their overexpression and mutant plants suggests that both of them are involved in Zn uptake and distribution within rice plants (Ishimaru et al., 2007; Lee et al., 2010b; Mu et al., 2021). In addition, higher expression level of *OsZIP4*, *OsZIP5*, *OsZIP8*, and *OsZIP10* was observed under Zn deficiency in the roots of accessions with the functional allele of OsHMA3, a tonoplast-localized transporter for Zn^2+^ and Cd^2+^ (Cai et al., 2019).

Relative to that in yeast and mammalian cells, little is known about sensing of Zn status in plant cells. Through yeast one-hybrid screening with the promoter sequence of Arabidopsis zinc deficiency-induced *AtZIP4* gene, two F-group bZIP transcription factors, AtbZIP19 and AtbZIP23, were identified to be involved in the adaptation of *Arabidopsis thaliana* to Zn deficiency (Assuncao et al., 2010). Although mutation of only one of them hardly affects plant growth and development, the *bzip19 bzip23* double mutant is hypersensitive to zinc deficiency (Assuncao et al., 2010). Orthologues of AtbZIP19 and AtbZIP23 are found in wheat, rice, poplar and even in the moss *Physcomitrella patens* (Assuncao et al., 2010; Evens et al., 2017; Lilay et al., 2020). These F-group bZIP transcription factors have a conserved N-terminal region that is enriched in cysteine and histidine residues, presumably to be a Zn-sensing domain (Lilay et al., 2019). Indeed, purified AtbZIP19 and AtbZIP23 could bind to Zn^2+^ in the size-exclusion chromatography coupled to inductively coupled plasma mass spectrometry (SEC–ICP-MS) assays *in vitro*, which is dependent on the cysteine- and histidine-rich (CHR) regions (Lilay et al., 2021), suggesting that AtbZIP19 and AtbZIP23 function as Zn-sensors in plants. Deletions or modifications of theses CHR domains of AtbZIP19/23 disrupt Zn^2+^ binding, leading to a constitutive transcriptional Zn deficiency response in Arabidopsis (Lilay et al., 2021). However, how CHR domains affect transcriptional activity of AtbZIP19/23 is not known. Based on phylogenetic analysis, OsbZIP48 (LOC_Os06g50310, also known as OsbZIP53) is the closest homolog of AtbZIP19 and AtbZIP23, while OsbZIP49 (LOC_Os01g58760, also known as OsbZIP07) and OsbZIP50 (LOC_Os05g41540, also known as OsbZIP44) are the closet homologs of AtbZIP24, another F-group bZIP transcription factor that is not involved in Zn deficiency response (Lilay et al., 2019; Lilay et al., 2020). Overexpression of *OsbZIP48* or *OsbZIP50*, but not *OsbZIP49*, complements the hypersensitive Zn deficiency phenotype of the Arabidopsis *bzip19 bzip23* double mutant, suggesting that OsbZIP48 and OsbZIP50 have a similar role to AtbZIP19 and AtbZIP23 in regulating Zn homeostasis (Lilay et al., 2020). However, the direct evidence for the role of OsbZIP48 and OsbZIP50 in Zn-sensing and Zn homeostasis in rice plants, and their downstream target genes under Zn deficiency await further investigation.

In the current study, we generate gene-edited mutant lines of *OsbZIP48* and *OsbZIP50*, and demonstrate that OsbZIP48 and OsbZIP50 regulate Zn/Fe/Cu homeostasis, especially under Zn deficiency in rice. We further report that the N-terminal CHR regions of OsbZIP48 and OsbZIP50 are important for their nucleus translocation. We also reveal the downstream genes of OsbZIP48 and OsbZIP50, and demonstrate that OsbZIP48 and OsbZIP50 directly bind to the promoter of a previous unidentified target gene, *OsZIP10*, and activate the ZDRE *cis*-element derived from *OsZIP10* promoter in effector-reporter assays. We further demonstrate the potential of editing *OsbZIP48* in the CHR domain for biofortification of Zn/Fe/Cu in brown rice. Thus, OsbZIP48 and OsbZIP50 are important for metal homeostasis under Zn deficiency in rice.

## Results

### Loss-of-function of *OsbZIP48* and *OsbZIP50* confers high sensitivity to Zn deficiency in rice

To explore the functional role of OsbZIP48/50 in Zn homeostasis in rice, we generated several loss-of-function lines of *OsbZIP48/50* double mutant with the CRISPR-Cas9 gene editing system (Figure S1A-B). Three double mutants, *bzip48-1 bzip50-1* (*dm-1*), *bzip48-2 bzip50-2* (*dm-2*), and *bzip48-3 bzip50-3* (*dm-3*) were characterized in details in the current study. In normal growth medium (0.4 μM Zn^2+^), shoot length and root length of *dm-1*/*dm-2/dm-3* were slightly reduced comparing to that of wild-type plants (ZH11), as a result, fresh weight and dry weight of *dm-1*/*dm-2/dm-3* mutants were also reduced (Figure 1A-B). Under Zn deficiency (no Zn^2+^ was added in the growth medium), the mutant phenotypes were severer than that in normal growth medium (Figure 1A-B). We also measured the content of Zn^2+^, Fe^2+^, Cu^2+^ in shoots and roots of these plants growing at different Zn levels, and found that the content of either Zn^2+^, Fe^2+^, or Cu^2+^ was obviously reduced in *dm-1*/*dm-2/dm-3* than that in ZH11 plants in shoots but less affected in roots (Figure 1C), suggesting that Zn/Fe/Cu translocations are impaired in *OsbZIP48/50* double mutant. To examine whether OsbZIP48/50 regulate plant responses to Fe deficiency, we grew ZH11 and *dm-1*/*dm-3* plants in Fe sufficient and Fe deficient growth medium. The results showed that there was no obvious difference between ZH11 and *dm-1*/*dm-3* plants under Fe deficiency in terms of shoot length and root length (Figure 1C). We also compared the phenotype between *OsbZIP48* single mutant and *OsbZIP48/50* double mutant, and found that the Zn deficiency-related phenotype was severer in *dm-1/dm-3* plants than that in *bzip48-1*/*bzip48-3* single mutant plants (Figure S2). Although we did not have the phenotypic data of *OsbZIP50* single mutant, these results suggest that OsbZIP48 and OsbZIP50 are functional redundant. We also generated *OsbZIP48-myc*/*OsbZIP50-myc* overexpression plants driven by the constitutive *Ubiquitin* promoter. While OsbZIP48-myc/OsbZIP50-myc fusion proteins were detected in transgenic rice plants by western blotting analysis (Figure S3-S4), Shoot height, root height, fresh weight and dry weight were not much different between these overexpression lines and wild-type plants under both Zn deficiency and sufficiency (Figure S3-S4). Together, these results demonstrate that OsbZIP48 and OsbZIP50 are required for Zn/Fe/Cu homeostasis under Zn deficiency in rice, and they regulate translocation of Zn/Fe/Cu from roots to shoots in rice.

**Figure 1.**
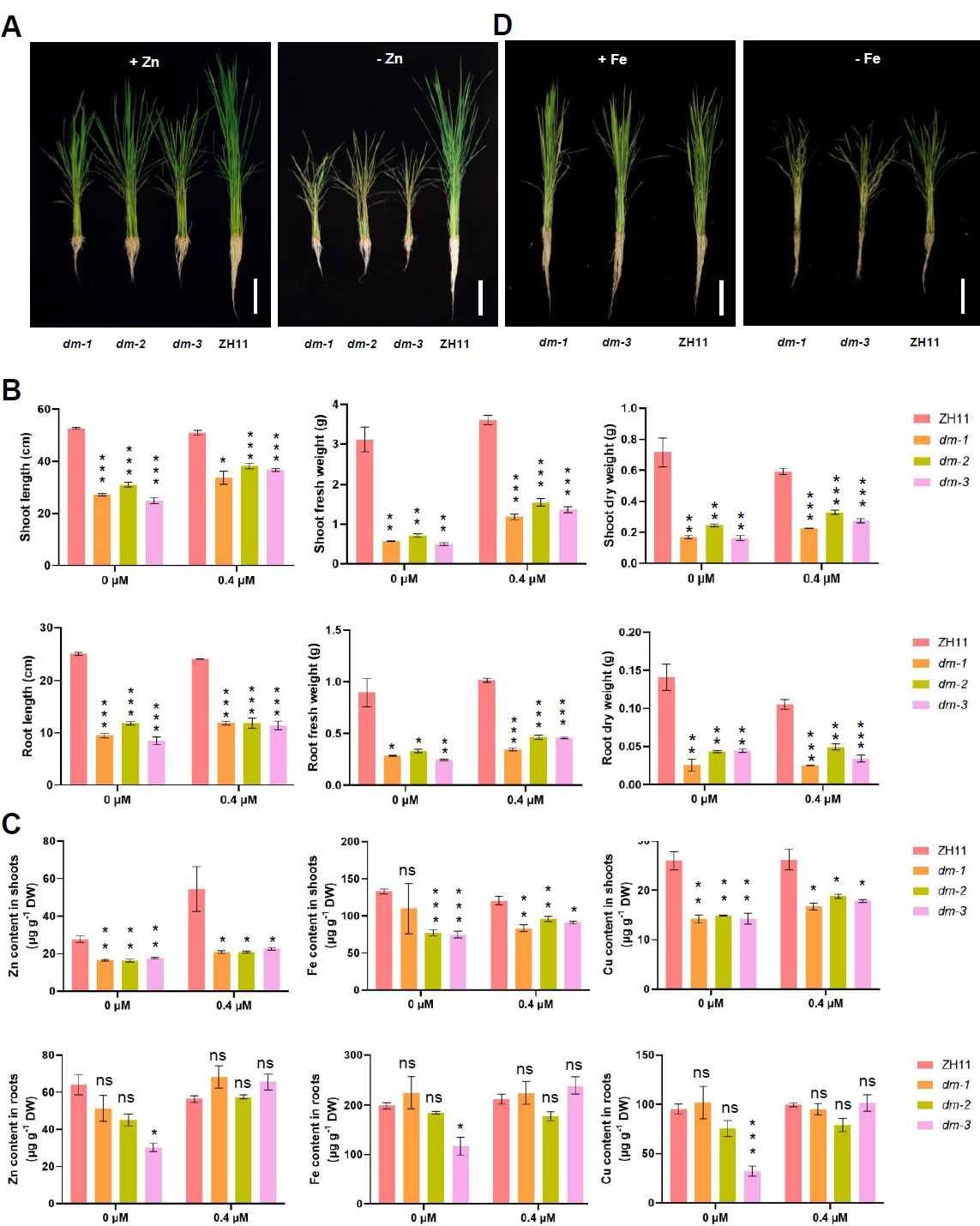
OsbZIP48 and OsbZIP50 Control Zn Homeostasis in Rice. (**A-B**) Phenotypic analysis of loss-of-function mutant of *OsbZIP48* and *OsbZIP50* grown in the presence (0.4 μM Zn^2+^) or absence of Zn. The double mutant of *OsbZIP48*/*50* (*dm-1*: *bzip48-1 bzip50-1*; *dm-2*: *bzip48-2 bzip50-2*; *dm-3*: *bzip48-3 bzip50-3*) and wild-type (ZH11) were grown in Kimura B medium for 3 weeks, and representative plants were photographed (A). Plant height, fresh weight and dry weight were measured (B). (**C**) Tissue elemental profiling determined with inductively coupled plasma-mass spectrometry (ICP-MS) in 3-week-old ZH11 and *dm-1*/*dm-1/dm-3* plants grown at different level of Zn supply. (**D**) Phenotypic analysis of loss-of-function mutant of *OsbZIP48* and *OsbZIP50* in the presence (20 μM Fe^2+^) or absence of Fe. ZH11 and *dm-1/dm-3* were grown in Kimura B medium in the presence of 40 μM Zn^2+^ for 3 weeks, and representative plants were photographed. Error bars represent SE (n=3 biological replicate). Asterisks indicate significance levels when comparing to ZH11 in t-test. (*, P<0.05; **, P<0.01; ***, P<0.001; ns, not significant at P<0.05). Bar = 10 cm in A and D.

### The CHR domain of OsbZIP48 and OsbZIP50 controls their subcellular localization

Both OsbZIP48 and OsbZIP50 have N-terminal CHR domains (Figure 2A), which presumably function as Zn-sensing domains (Lilay et al., 2021). When OsbZIP48 or OsbZIP50 was expressed in Arabidopsis *bzip19 bzip23* double mutant, they were found in both cytosol and nucleus (Lilay et al., 2020), but how these different subcellular localization is regulated remains unexplored. We expressed the YFP-tagged form of full-length or truncated forms (Figure 2A) of OsbZIP48/50 in tobacco epidermal cells, and found that the full-length OsbZIP48-YFP and OsbZIP50-YFP were localized both in cytosol and nucleus, and OsbZIP48-YFP was predominantly localized in cytosol while OsbZIP50 was predominantly localized in nucleus (Figure 2B-C). When one or two CHR domains at N-terminus of OsbZIP48/50 were removed, the truncated proteins were mainly observed in nucleus (Figure 2B-C). These results suggest that the CHR domain of OsbZIP48/50 is important for their nucleus localization. To further confirm these results, we mutated the cysteine and histidine amino acids in one or two CHR domains of OsbZIP48/50 (Figure 3A), and expressed the YFP fusion proteins in tobacco epidermal cells. The results showed that different from the native OsbZIP48-YFP or OsbZIP50-YFP, the mutated forms of OsbZIP48/50 in CHR domains (bZIP48m1/50m1 or bZIP48m2/50m2 or bZIP48m3/50m3) were observed solely in nucleus (Figure 3B-C). These results further support that the CHR domain of OsbZIP48/50 regulates their nucleus localization, therefore, further transcriptional activity.

**Figure 2.**
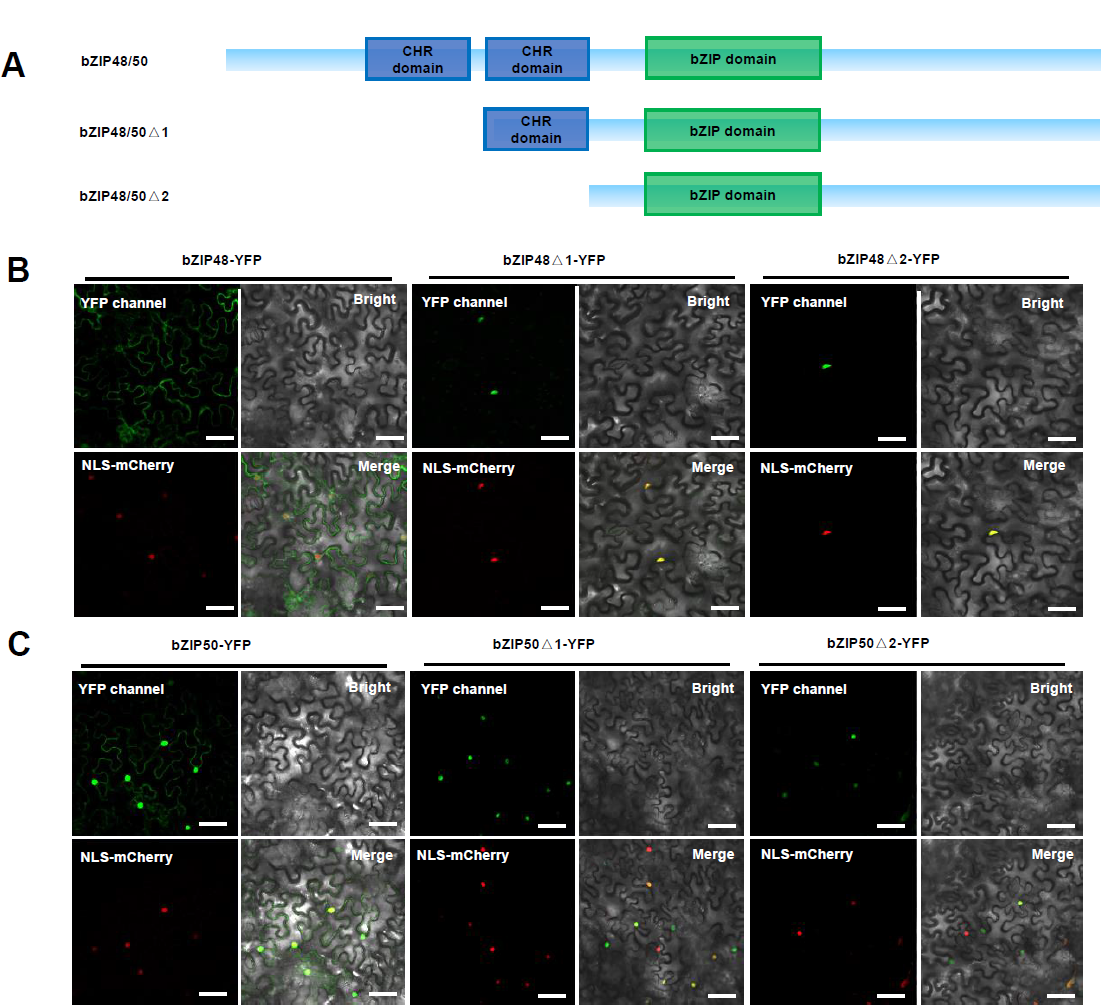
The N-terminus of OsbZIP48 Is Important for Its Nucleus Localization. (**A**) Diagram showing the full-length and different truncations of OsbZIP48/50. CHR domain: cysteine- and histidine-rich domain; bZIP domain: basic leucine zipper domain. (**B-C**) Subcellular localization of the full-length and two truncated forms of OsbZIP48/50 in tobacco epidermal cells. NLS-mCherry was used as a nucleus marker. Bar = 50 μm.<colcnt=3>

**Figure 3.**
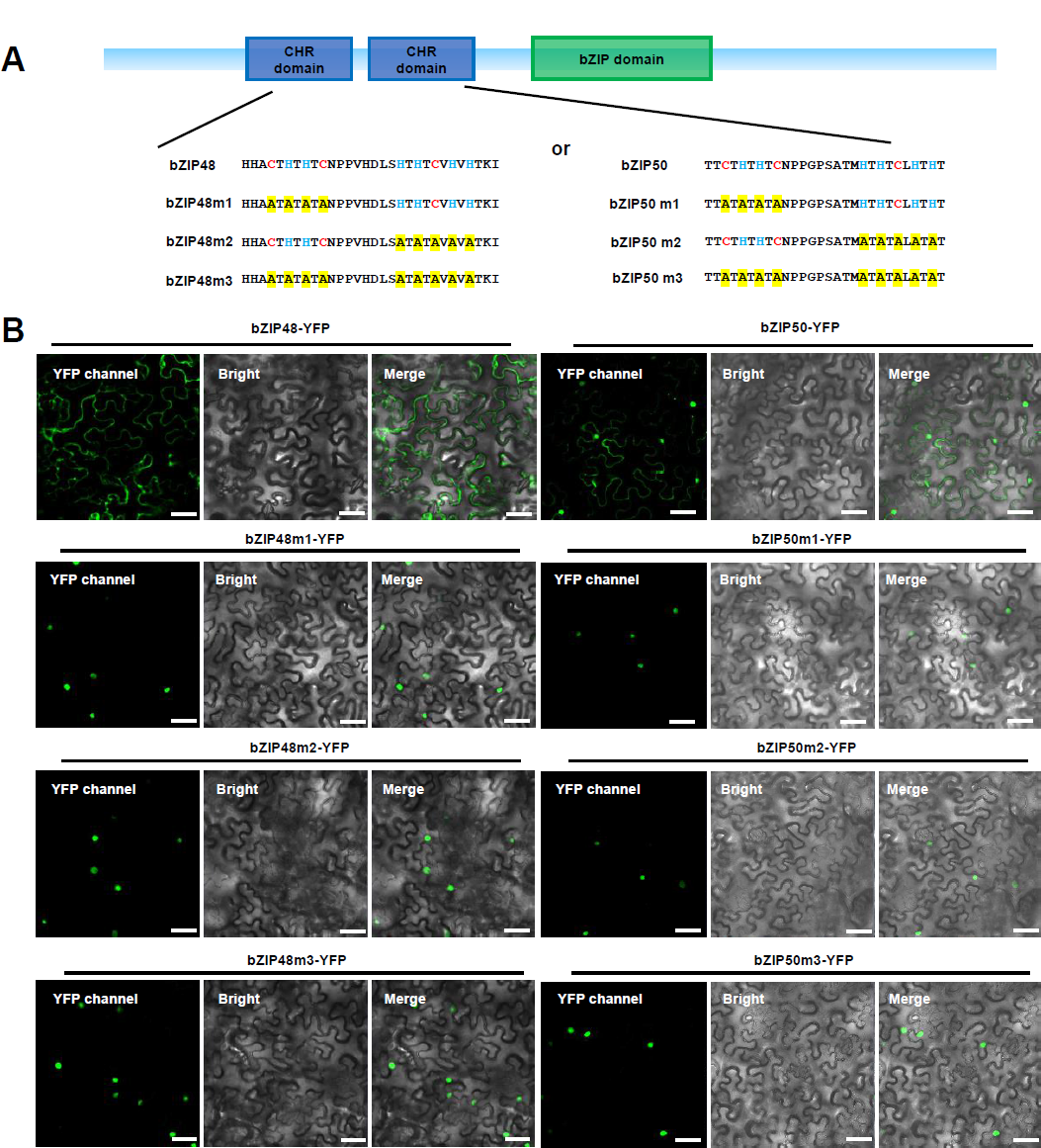
The CHR Domains of OsbZIP48/50 Are Important for Their Nucleus Localizations. (**A**) Diagram showing the cysteine- and histidine-rich (CHR) domain of native and mutated form of OsbZIP48/50. (**B-C**) Subcellular localization of the native and three mutated forms of OsbZIP48/50-YFP in tobacco epidermal cells. Bar = 50 μm.

### OsbZIP48 and OsbZIP50 regulate Zn-deficit responsive genes in rice

To examine whether OsbZIP48/50 regulate Zn-deficit responsive gene expression, the transcriptional activation activity of OsbZIP48/50 was firstly tested in yeast cells in which the full-length of OsbZIP48 or OsbZIP50 was fused to GAL4 DNA-binding domain, respectively. Compared to the empty vector control (BD), yeast cells expressing the fusion proteins (BD-bZIP48 or BD-bZIP50) grew well on synthetic droplet mediums (Figure 4A). These results suggest that OsbZIP48/50 have transcriptional activation activity. Then we performed RNA-Seq analysis with ZH11 and *OsbZIP48/50* double mutant (*dm-1*) grown under Zn-sufficient and Zn-deficient conditions. In ZH11 plants, totally 1177 and 469 genes were significantly up-regulated (log_2_FC ≥ 1, q ≤ 0.05) and down-regulated (log_2_FC ≤ −1, q ≤ 0.05), respectively (Figure 4B and **Supplemental Data 1**). Among them, 1117 genes were not significantly up-regulated (log_2_FC ≥ 1, q ≤ 0.05) and 278 genes were not significantly down-regulated (log_2_FC ≤ −1, q ≤ 0.05) in *dm-1* mutant plants, respectively (Figure 4B and **Supplemental Data 1**). Since OsbZIP48/50 have transcriptional activation activity (Figure 4A), we considered these 1117 genes as *OsbZIP48/50*-dependent downstream genes. Gene Ontology (GO) enrichment analysis of these 1117 genes was further carried out and the results showed that phenylpropanoid biosynthetic process, peroxidase activity, fatty acid biosynthetic process, microtubule, cell wall organization, et al., were enriched (Figure 4C). We selected 12 genes including *OsZIP4* and *OsZIP10* in the ZIP transporter family, and genes involved in phenylpropanoid biosynthesis, and performed RT-qPCR to validate the RNA-Seq results. All these 12 genes including *OsZIP4* and *OsZIP10* were significantly up-regulated by Zn-deficiency in ZH11, but they were not much induced by Zn-deficiency in *dm-1* plants (Figure 5A). Thus, OsbZIP48 and OsbZIP50 are important for Zn-deficit responsive gene expression in rice.

**Figure 4.**
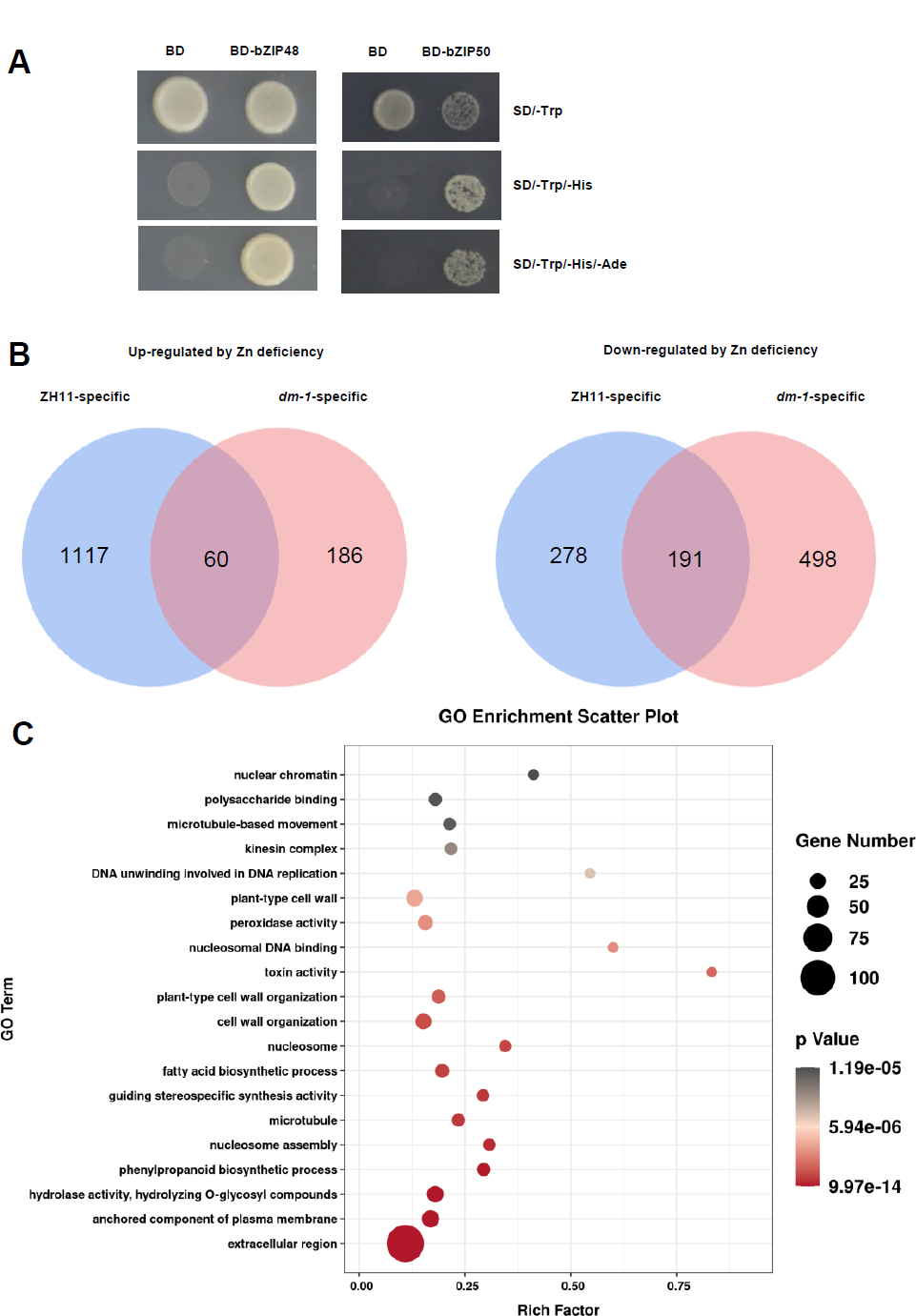
OsbZIP48 and OsbZIP50 Regulate Global Gene Expression under Zn Deficiency in Rice. (**A**) Transcriptional activation activity of OsbZIP48/50. The full-length of OsbZIP48 or OsbZIP50 was fused to the yeast GAL4 DNA-binding domain (BD) and activation of the *His* and *Ade* reporter genes in yeast cells was checked in the droplet growth medium (minimal synthetic defined (SD) bases with droplet supplement minus Trp, or minus Trp and His or minus Trp, His, and Ade). (**B**) Venn diagrams showing the numbers of overlapping and non-overlapping genes that were up-regulated or down-regulated in wild-type (ZH11) and *OsbZIP48/50* double mutant (*dm-1, bzip48-1 bzip50-1*) grown under Zn sufficiency and deficiency. Criteria for differential expression was set as q ≤ 0.05, fold change (FC) ≥ 2 for up-regulation or FC ≤ 0.5 for down-regulation. (**C**) GO analysis of 1117 genes that were specifically up-regulated in ZH11 by Zn deficiency.

**Figure 5.**
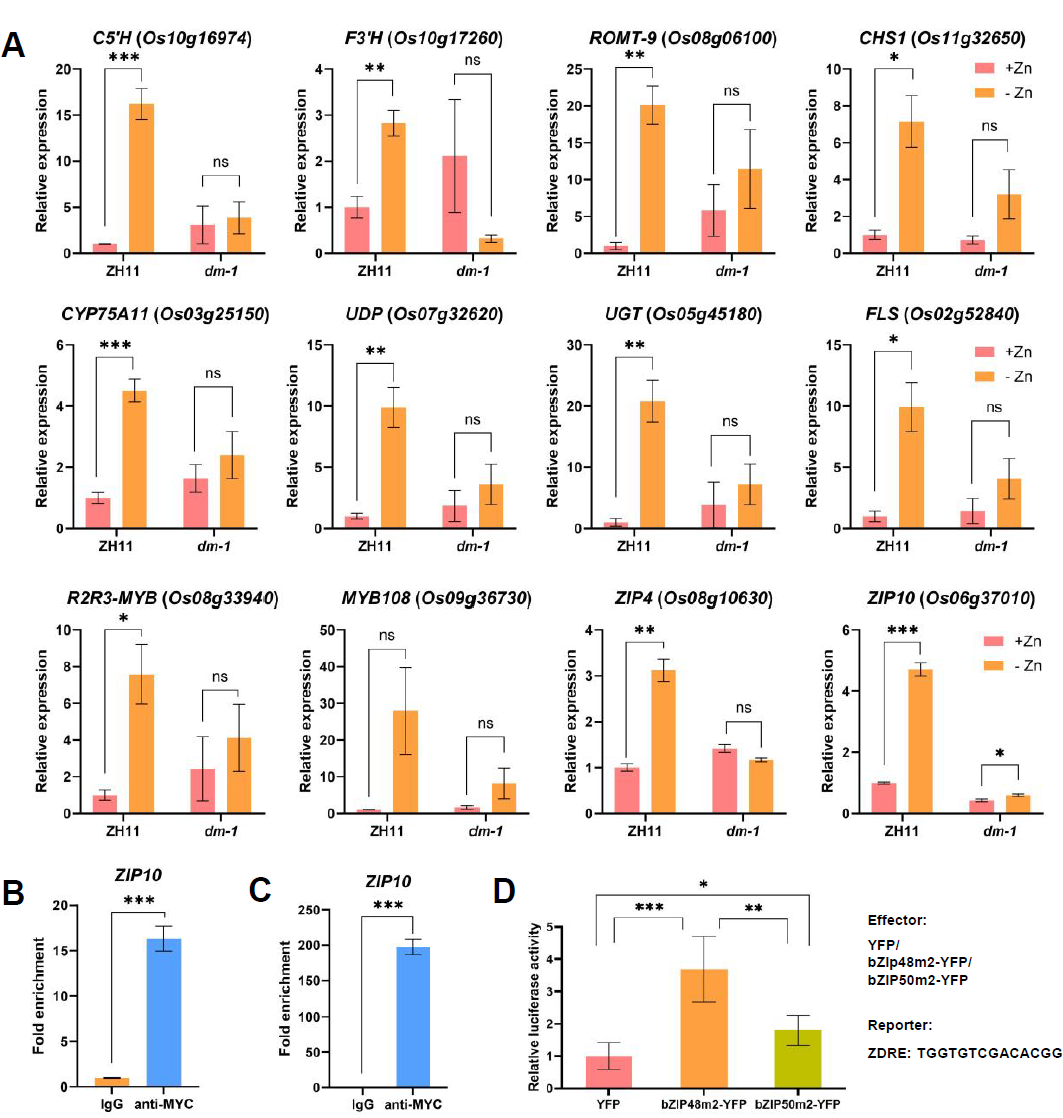
OsbZIP48 and OsbZIP50 Directly Regulate the Expression of *OsZIP10* in Rice. (**A**) Validation of OsbZIP48/50 downstream gene expression by RT-qPCR. Wild-type (ZH11) and *OsbZIP48/50* double mutant (*dm-1, bzip48-1 bzip50-1*) grown under Zn sufficiency and deficiency were harvest for RT-qPCR. Relative gene expression is the expression level of target gene under Zn deficiency relative to that under Zn sufficiency, both of which were normalized to that of the internal control *ACTIN*. (**B-C**) Direct binding of OsbZIP48/50 to the *OsZIP10* promoter in rice. Transgenic plants expressing *OsbZIP48-myc* (B) or *OsbZIP50-myc* (C) in ZH11 background were harvested for ChIP-qPCR analysis using anti-myc antibody. IgG was used as a negative control. Fold enrichment of each sample was normalized to that of IgG sample, both of which were normalized to that of the *UBQ5* control. (**D**) Activation of ZDRE *cis*- element by OsbZIP48/50 in effector-reporter assays. YFP fusion with the mutated form of OsbZIP48/50 driven by the CaMV 35S promoter was expressed as the effector, and the promoter region of *OsZIP10* containing ZDRE was linked to the firefly luciferase and used as a reporter. Renilla luciferase driven by the CaMV 35S promoter was used as an internal control. Relative luciferase activity is the firefly luciferase activity normalized to the Renilla luciferase activity, which was then normalized to the empty vector control (YFP). The second CHR domain of OsbZIP48/50 was mutated. Error bars represent SE (n=3 biological replicate) in A-C. Error bars represent SD (n=5) in D. Asterisks indicate significance levels in t-test. (*, P<0.05; **, P<0.01; ***, P<0.001; ns, not significant at P<0.05).

### OsbZIP48 and OsbZIP50 directly regulate the expression of *OsZIP10*

OsZIP10 is predicted to be involved in divalent cation (Zn^2+^, Fe^2+^, and Cu^2+^) homeostasis in plants (Grotz, et al., 1998). To examine whether OsbZIP48/50 directly control the expression of *OsZIP10* under Zn deficiency, we performed Chromatin Immunoprecipitation-quantitative PCR (ChIP-qPCR) analysis using *OsbZIP48-myc/OsbZIP50-myc* overexpression plants. The results showed that *OsZIP10* promoter DNA was highly enriched when *anti*-myc antibody was used in the ChIP-qPCR assays (Figure 5B-C). We concluded that OsbZIP48/50 directly bind to the promoter of *OsZIP10*. Previously, specific Zinc-Deficiency Response Element (ZDRE: RTGTCGACAY) motif was found in the promoter region of AtbZIP19/23-regulated genes (Lilay et al., 2020). We checked the promoter region of *OsZIP10*, and found that there are three copies of ZDRE motifs presented in 1 kb promoter region of *OsZIP10*. We linked the promoter region of *OsZIP10* containing ZDRE to firefly luciferase reporter, and performed dual-luciferase reporter assays using the nucleus-localized form of OsbZIP48 (OsbZIP48m2-YFP) or OsbZIP50 (OsbZIP48m2-YFP) as effector in tobacco leaves, the relative luciferase activity was higher when OsbZIP48m2-YFP or OsbZIP50m2-YFP was co-infiltrated with the reporter than that when empty vector control (YFP) was co-transformed (Figure 5D). Taken together, these results suggest that OsbZIP48 and OsbZIP50 directly regulate the expression of *OsZIP10* under Zn deficiency in rice.

### Targeted gene-editing of *OsbZIP48* increases Zn/Fe/Cu content in brown rice

Because the CHR domain of OsbZIP48 inhibits its nucleus localization (Figure 2 & Figure 3), we were interested in whether manipulation of this locus by gene editing could improve micronutrients in rice grains. We obtained two double mutants (*dm-4*: *bzip48-4 bzip50-3* and *dm-5*: *bzip48-5 bzip50-4*) in which the CHR domain of OsbZIP48 is disrupted and the bZIP domain of OsbZIP50 is absent (Figure S1). These double mutants were grown together with ZH11 in Changxing Experimental Station of Zhejiang University (Huzhou, Zhejiang) with standard cultivation conditions. At maturity, the content of Zn/Fe/Cu was measured in flag leaves and brown rice seeds. Comparing to the ZH11 control plants, the content of Zn/Fe/Cu of *dm-4/dm-5* plants was much higher in both flag leaves and brown rice (Figure 6). Thus, these preliminary results showed the potential of editing *OsbZIP48* for improving micronutrients in rice grains.

**Figure 6.**
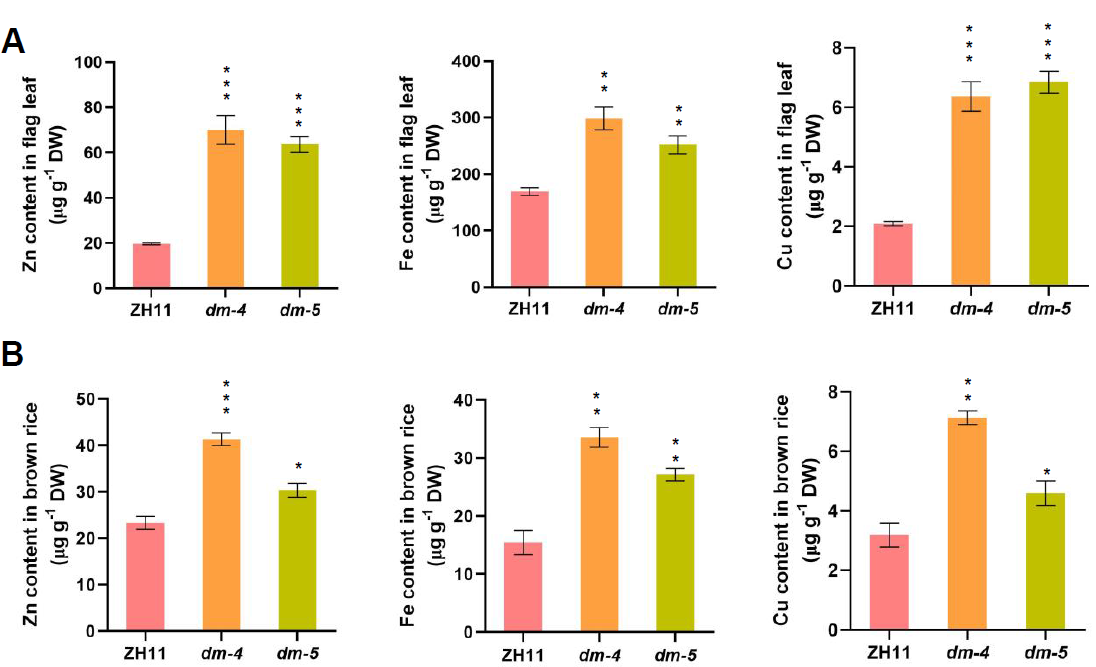
Mutation of OsbZIP48 CHR domain in *OsbZIP50* mutant background increases Zn/Fe/Cu content in brown rice. (**A-B**) Tissue elemental profiling determined with inductively coupled plasma-mass spectrometry (ICP-MS) in flag leaves (A) and brown rice (B) of ZH11 and *dm-4*/*dm-5* plants grown in the field. Error bars represent SE (n=3 biological replicate). Asterisks indicate significance levels when comparing to ZH11 in t-test (*, P<0.05; **, P<0.01; ***, P<0.001).

## Discussions

Zn is one of the most important essential micronutrients, and thus required for normal growth and development of plants. Zn homeostasis in plants is a complexed regulatory network including Zn uptake, accumulation, trafficking, sequestration, remobilization, redistribution, and detoxification (Ajeesh Krishna et al., 2020). Although previous preliminary results showed that OsbZIP48 and OsbZIP50 have a conserved function related to Zn homeostasis in Arabidopsis (Lilay et al., 2020), our current results provide direct evidence on the important role of OsbZIP48/50 in Zn/Fe/Cu homeostasis in rice, since loss-of-function mutation of *OsbZIP48/50* greatly impairs plant growth and metal accumulation, especially under Zn deficiency. However, overexpression of *OsbZIP48* or *OsbZIP50* could not improve plant growth under Zn deficiency, suggesting that the function of OsbZIP48/50 in Zn homeostasis in rice requires other additional unknown factors. Identification of OsbZIP48/50 interacting factors will enhance the understanding of regulatory mechanism controlled by OsbZIP48/50.

Plant ZIP family proteins are membrane transporters involved in cellular uptake of Zn, but they also transport other divalent metal cations such as Fe^2+^ and Cu^2+^ (Ajeesh Krishna et al., 2020). In the current study, the expression of *OsZIP4* and *OsZIP10* is impaired in *OsbZIP48/50* double mutant, especially under Zn deficiency, and OsbZIP48/50 could activate the promoter activity of *OsZIP10*, which is agreed with the results that the content of Fe^2+^ and Cu^2+^ is also reduced in shoots of *OsbZIP48/50* double mutant compared to that in wild-type ZH11 plants. However, the sensitivity of *OsbZIP48/50* double mutant to Fe deficiency is not much affected, suggesting that the amount of Fe^2+^ adsorbed by other types of transporters such as IRT1 (Dubeaux et al., 2018) is sufficient in *OsbZIP48/50* double mutant, and OsbZIP48/50 play a minor role in response to Fe deficiency. Previous results showed that OsZIP4 is involved in Zn uptake and redistribution, the function of OsZIP10 in Zn uptake and translocation needs further investigated. Nevertheless, our results demonstrate that OsbZIP48/50 are important not only Zn homeostasis, but also for Fe and Cu homeostasis under Zn deficiency.

The Zn sensing mechanism in plants is little understood. In yeast (*Saccharomyces cerevisiae*), the Zinc-dependent activator protein-1 (Zap1) encodes a transcriptional activator with multiple Cys(2)His(2) zinc fingers and functions as a Zn sensor (Bird et al., 2003). Zap1 binds to Zn^2+^ through two zinc fingers in the activation domains, which represses its transcription activity in Zn replete cells, while in Zn-limited cells, Zap1 is activated and induces the expression of genes involved in Zn uptake (Bird et al., 2003). Similar mechanism has been recently revealed in Arabidopsis, in which two bZIP transcription factors AtbZIP19 and AtbZIP23 serve as Zn sensors (Lilay et al., 2021). Under Zn sufficient conditions, binding Zn^2+^ to AtbZIP19/23 in the CHR region inhibits the transcriptional activity of AtbZIP19/23, while under Zn deficient conditions, AtbZIP19/23 are activated and induce downstream gene expression (Lilay et al., 2021). In the current study, we further demonstrate that the CHR region of OsbZIP48/50 is important for their nucleus localization (Figure 7). Ectopic expression of a mutated form of AtbZIP19 in the Zn-sensing domain in *AtbZIP19/23* double mutant background results in constitutively transcriptional Zn deficiency response in Arabidopsis (Lilay et al., 2021). We further demonstrate that deletion one or 19 amino acids in the CHR domain of OsbZIP48 in *OsbZIP50* mutant background increases Zn/Fe/Cu content in brown rice. Therefore, modification of these Zn-sensing domains in OsbZIP48/50 by gene-editing especially through prime-editing-library screening (Xu et al., 2021) could provide a possible strategy for breeding Zn-efficient and biofortified rice cultivars in future.

**Figure 7.**
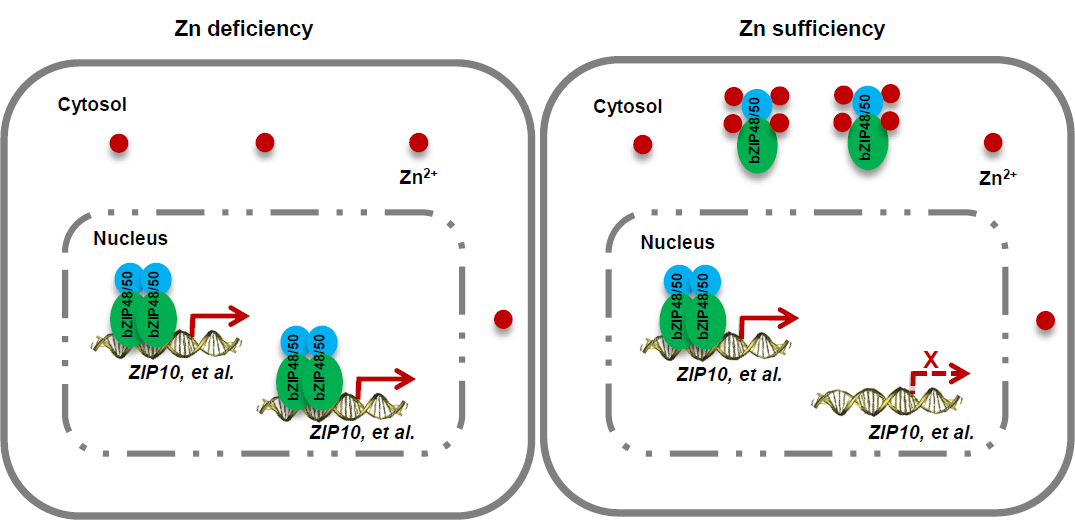
A working model for the role of OsbZIP48 and OsbZIP50 in Zn homeostasis in rice. Under Zn sufficiency, OsbZIP48/50 are retained in cytosol due to the excess of Zn^2+^. Under Zn deficiency, OsbZIP48/50 are disassociated with Zn^2+^ and have some configuration changes, which leads to predominantly nucleus relocation. OsbZIP48/50 recognize ZDRE *cis*-element presented in the promoter of downstream genes such as *OsZIP10* in nucleus and activate their expression under Zn deficiency. These downstream genes are important for Zn translocation in rice.

## Materials and methods

### Plant materials and growth conditions

In this study, all the gene-editing mutants were in ZH11 background, while the overexpression materials were in Nipponbare background. Rice seeds were soaked for two days at 37°C and then transferred to 96-well plates containing Kimura B nutrient solution to grow under the following conditions: 20,000 lux white fluorescent light (12 h light/ 12 h dark) at 29°C and with 65% relative humidity. For Zn-deficiency phenotype experiments, germinated seeds were transferred to Kimura B nutrient solutions supplemented with different concentrations of Zn^2+^ for three weeks and then photographed. Plant height, root length, fresh weight, dry weight was then measured, respectively. For Fe-deficiency experiments, plants were grown in Kimura B nutrient solution supplemented with different concentrations of Fe in the presence of 40 μM Zn^2+^. All nutrient solutions used in the experiment were prepared using distilled water, and 12 plants in three biological replicates were phenotypically analysed.

### Targeted gene editing and transgenic overexpression

To generate gene-edited plants using the CRISPR-Cas9 technology, gene-specific guide sequences (sgRNAs) were designed, and the expression cassettes were cloned into the CRISPR-Cas9 system, To overexpress the full-length myc-tagged OsbZIP48 or OsbZIP50, the coding sequence of *OsbZIP48* or *OsbZIP50* and myc tag were amplified with PCR and inserted into pCAMBIA1301 to produce ProUbiqutin::OsbZIP48-myc or ProUbiqutin::OsbZIP50-myc expression constructs. Error-free constructs were introduced into plants by Agrobacterium-mediated stable transformation, and transgenic-free mutants and overexpression transgenic plants were screened and maintained.

### Elemental analysis of plant tissues

To determine the elemental content, plant tissues were washed with 20 mM disodium EDTA for 15 to 20 minutes, followed by three washes with 5 mM CaCl_2_ and twice in ultrapure water. The tissues were then blotted dry with absorbent paper and dried to a constant weight at 65°C. For sample preparation, 0.1 g of homogenized sample was taken and digested with HNO3-H_2_O_2_ (5:1 [v/v]) acid mixture. Elemental content was determined by inductively coupled plasma mass spectrometry (ICP-MS) .

### Microscopy analysis

For subcellular localization study, the YFP tag was fused to the full-length, various truncated and mutated form of OsbZIP48 and OsbZIP50, respectively. Error-free constructs were transiently expressed in tobacco (*Nicotiana benthamiana*) leaves via Agrobacterium-mediated transformation. Samples were taken and prepared for observation using confocal microscopy (Zeiss LSM A710). A nuclear localization signal fused with mCherry (NLS-mCherry) was used as a nucleus marker.

### RNA-seq and RT–qPCR analysis

For RNA-seq analysis, rice seedlings grown in Kimura B nutrient solutions containing either 0 μM or 0.4 μM Zn ions for 5 days with three biological replicates were collected and immediately froze in liquid nitrogen. Total RNA was extracted with Trizol reagent (Thermofisher, USA) following the manufacturer’s procedure. The cDNA libraries were constructed following Illumina standard protocols and sequenced on the Illumina Novaseq™ 6000 (LC-Bio Technology CO., Ltd., Hangzhou, China). The RNA-seq reads were aligned to the MSU rice reference genome using Hisat2. The gene expression level was calculated and normalized to FPKM (fragments per kilobase of transcript per million mapped reads) with Stringtie. Differential expression analysis of genes was performed by DESeq2 software between two different groups. The genes with the parameter of false discovery rate (FDR) below 0.05 and absolute fold change ≥ 2 were considered as differentially expressed genes. For RT-qPCR, total RNA was extracted using a RAN Prep Pure Plant Kit (Tiangen, Beijing, China). For reverse transcription, 1000ng of RNA was reverse transcribed into cDNA in a 20 μL reaction using the Evo M-MLV RT Master Mix (AG, Changsha, China). Quantitative RT-PCR was performed using the SuperReal PreMix Color (Tiangen, Beijing, China) in the CFX96 real-time system (Bio-Rad, Hercules, CA, USA). The gene expression levels in three biological replicates were calculated using the ΔΔCt (threshold cycle) method. *ACTIN* was used as an internal control for normalization.

### ChIP-qPCR

Chromatin Immunoprecipitation (ChIP) was performed according to standard protocols. One-week-old *OsbZIP48-myc*/*OsbZIP50-myc* transgenic seedlings grown at Kimura B nutrient solution supplied with 0.4 μM Zn^2+^ were transfer to Kimura B nutrient solutions without Zn^2+^ for another one week and then collected for ChIP assays. Protein G agarose (Millipore, Temecula, CA, USA) and anti-myc antibody (Abmart, Shanghai, China) or IgG (Abmart) were used for immunoprecipitation. The ChIP products were analysed by quantitative PCR.

### Transcriptional activation activity assays

To examine the transactivation activity of OsbZIP48 or OsbZIP50, the coding sequence (CDS) of *OsbZIP48* or *OsbZIP50* was amplified from ZH11, cloned into the pGBKT7 vector, and transformed into yeast strain P69J-4A. The transformed yeast strains were examined on SD/-Trp, SD/-Trp/-His and SD/-Trp/-His/-Ade plates for 3-4 days at 30°C.

### Effector-reporter assays

For the LUC activity assays, 3 copies of ZDRE sequence (TGGTGTCGACACGG) derived from the promoter region of *OsZIP10* was inserted into pGreen0800-II with the 35S minimal promoter included to generate the reporter vector. The mutated form of OsbZIP48 or OsbZIP50 (OsbZIP48m2 or OsbZIP50m2) fused with YFP were inserted into the expression vector as effectors. Error-free constructs were transiently expressed in tobacco (*Nicotiana benthamiana*) leaves via *Agrobacterium tumefaciens* strain GV3101. The infiltrated plants were grown for an additional 3 days, and luciferase activities were measured using a dual-luciferase reporter assay kit (Promega, Beijing China).

### Western blotting analysis

For Western blotting analysis, total proteins were extracted with the extraction buffer (125 mM Tris-HCl, pH 8.0, 375 mM NaCl, 2.5 mM EDTA, 1% SDS and 1% β-mercaptoethanol) from the transgenic rice plants. After that, proteins were separated in 4%-10% SDS-PAGE gels and analysed using anti-myc antibody (Abmart). Anti-Histone (H3) was used as a loading control.

## Data availability

The RNA-Seq data is deposited in the Genome Sequence Archive (GSA) under the accession number CRA010766.

## Supplementary materials

Supplementary material for this article is available at XXXXXXXX

## Acknowledgements

This project was financially supported by grants from the State Key Project of Research and Development Plan (2021YFF1000404), and Natural Science Foundation of Zhejiang, China (LD21C020001).

## Author contributions

J.X.L. and T.Q. designed the experiments; T. Q., T. C.X. Q.Y.Z. and H.P.L. performed the experiments; J.X.L. and T.Q. analyzed the data; J.X.L., and T.Q. wrote the paper.

## Declaration of interests

The authors declare no competing interests.

## Supplemental Materials

**Figure S1.**
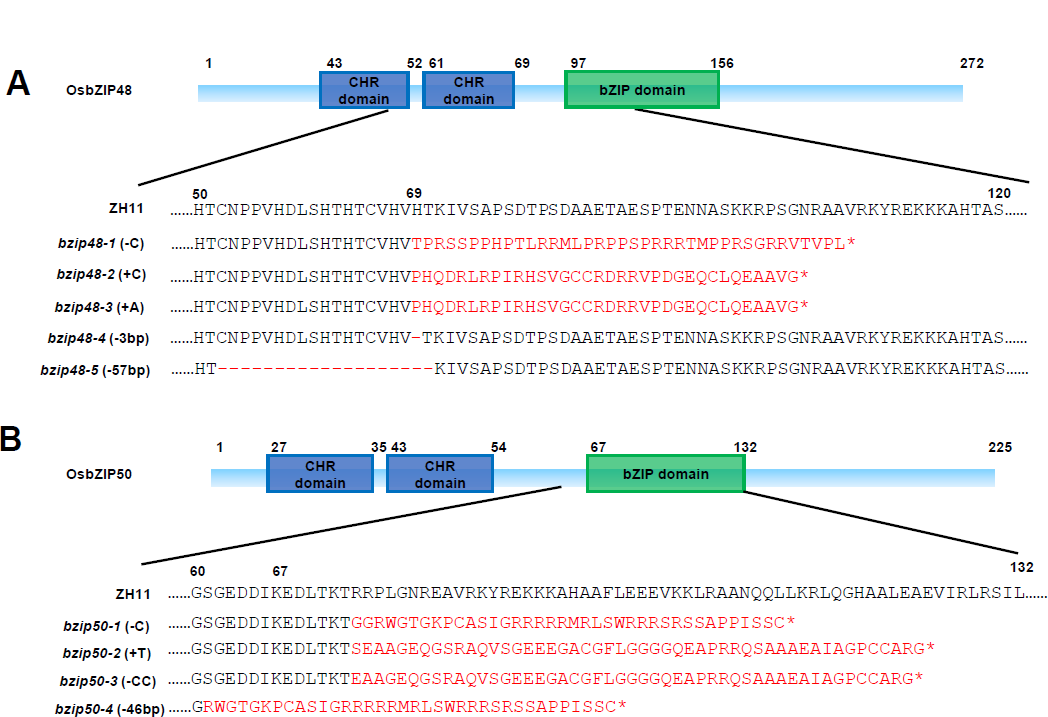
Characterization of Os*bZIP48/50* Mutants. (**A-B**) Partial protein sequences of OsbZIP48/50 in wild-type (ZH11) and gene-edited mutants of OsbZIP48 (A) or OsbZIP50 (B) are aligned. The newly derived amino acids in mutants are shown in red. CHR domain: cysteine- and histidine-rich domain; bZIP domain: basic leucine zipper domain. *, stop codon.

**Figure S2.**
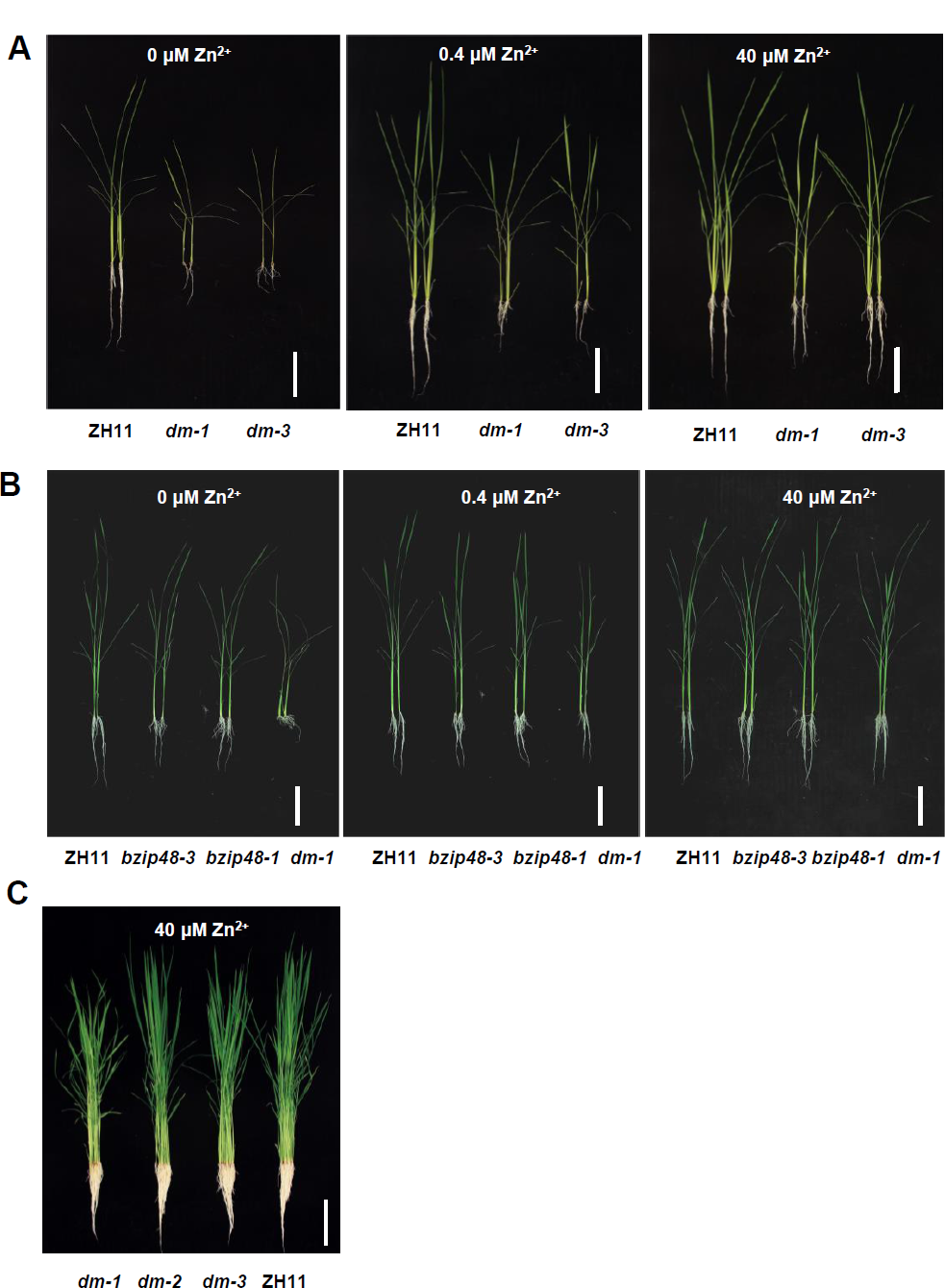
Comparison of the Phenotype between *OsbZIP48* Single Mutant and *OsbZIP48/50* Double Mutant. (**A-C**) Phenotypic analysis. The single mutants of OsbZIP48 (*bzip48-1*/*bzip48-3*), double mutant of *OsbZIP48*/*50* (*dm-1, bzip48-1 bzip50-1*; *dm-2, bzip48-2 bzip50-2*; *dm-3, bzip48-3 bzip50-3*), and wild-type (ZH11) were grown in Kimura B medium supplied with different concentration of Zn^2+^ for 21 days (A, C) or 18 days (B), and representative plants were photographed. Bar = 10 cm.

**Figure S3.**
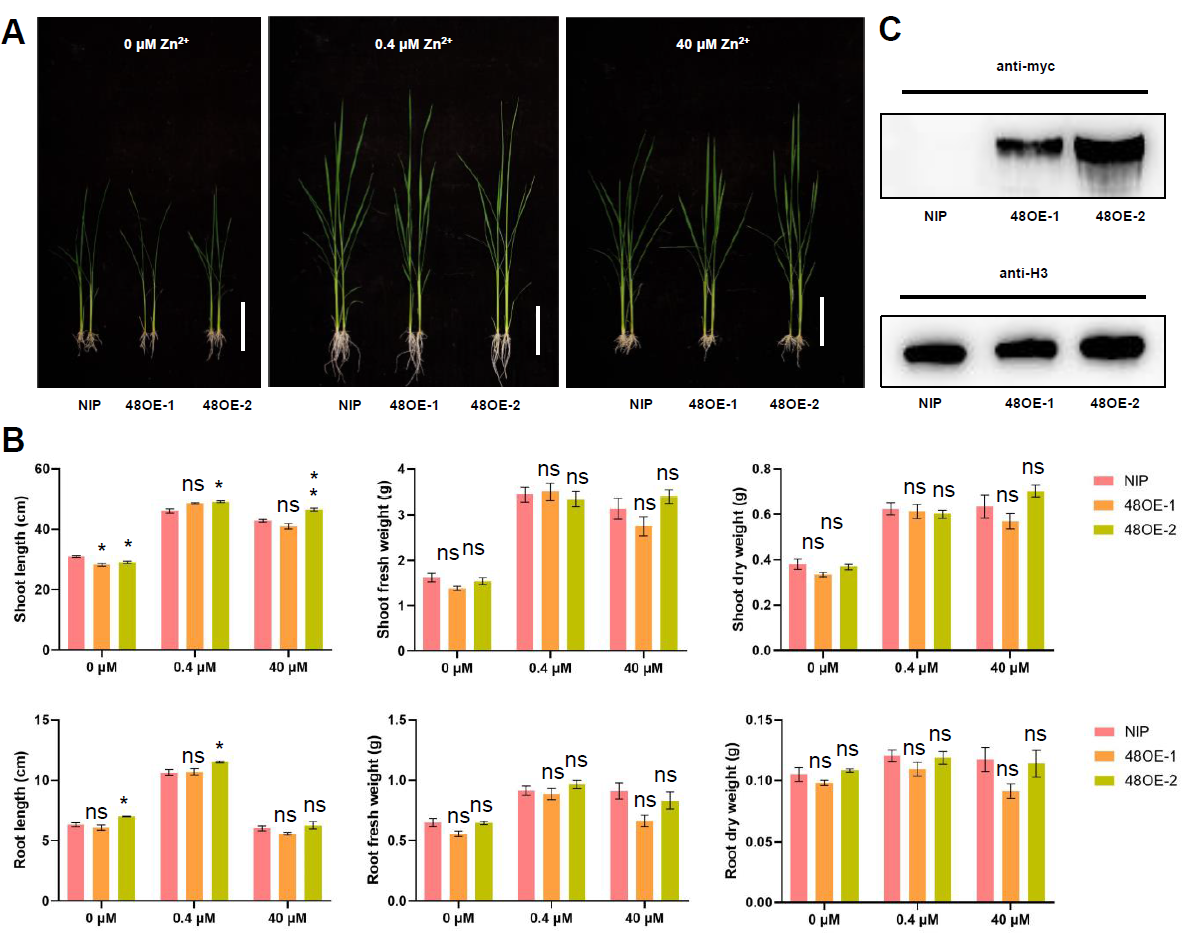
Phenotypic analysis of OsbZIP48 Overexpression Plants. (**A-B**) Phenotypic analysis. The OsbZIP48-myc overexpression (OE48-1/OE48-2) and wild-type (NIP) plants were grown in Kimura B medium supplied with different concentration of Zn^2+^ for 3 weeks, and representative plants were photographed (A). Plant height, fresh weight and dry weight were measured (B). The expression of level of transgene at 0.4 μM Zn^2+^ was validated by Western blotting analysis using anti-myc antibody, anti-His antibody was used as an internal loading control. Bar = 10 cm.

**Figure S4.**
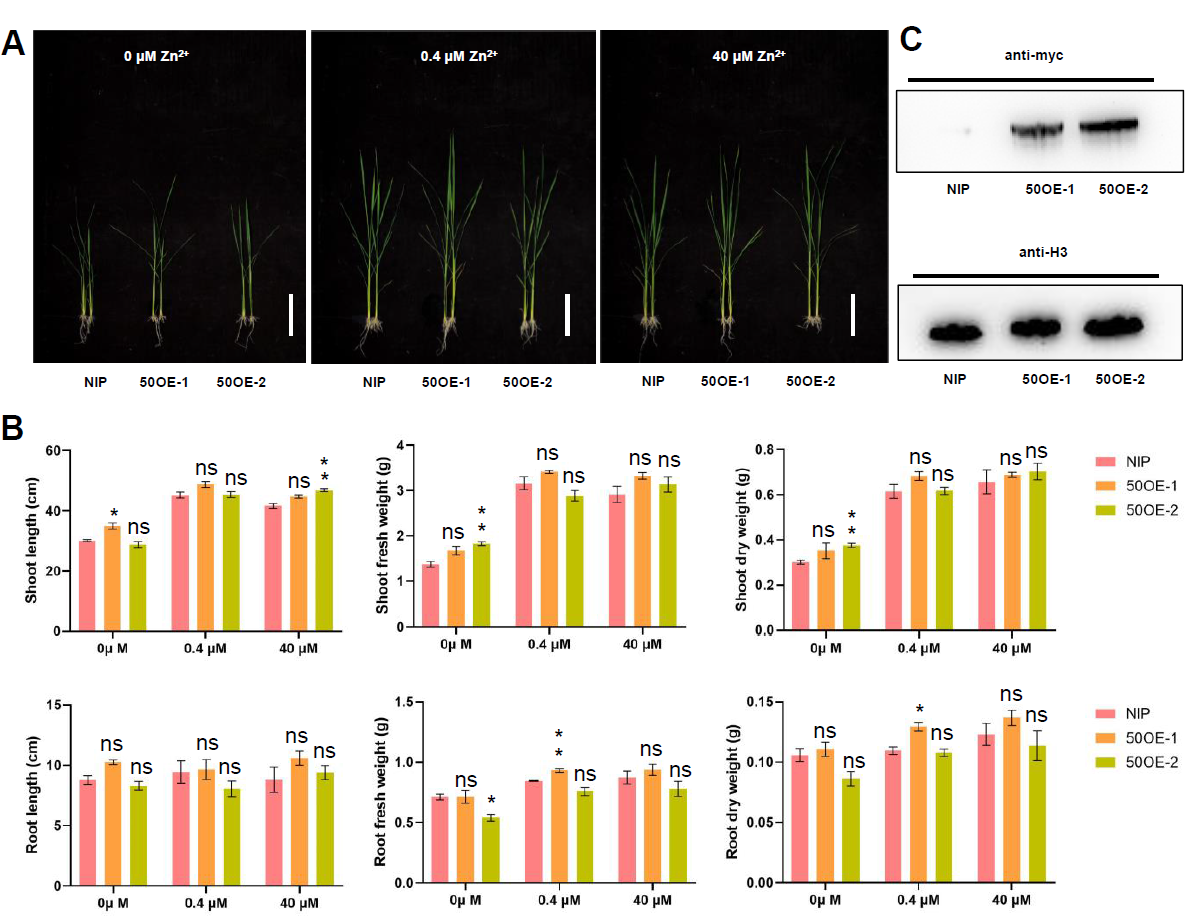
Phenotypic analysis of OsbZIP50 Overexpression Plants. (**A-B**) Phenotypic analysis. The OsbZIP50-myc overexpression (OE48-1/OE48-2) and wild-type (NIP) plants were grown in Kimura B medium supplied with different concentration of Zn^2+^ for 3 weeks, and representative plants were photographed (A). Plant height, fresh weight and dry weight were measured (B). The expression of level of transgene at 0.4 μM Zn^2+^ was validated by Western blotting analysis using anti-myc antibody, anti-His antibody was used as an internal loading control. Bar = 10 cm.

**Figure S5.**
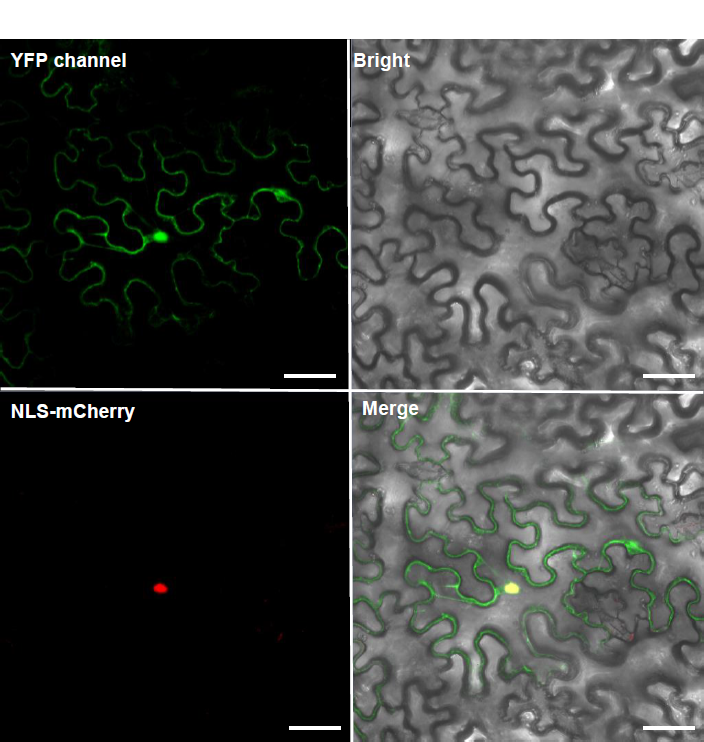
Subcellular Localization of the YFP Control. The empty vector (YFP) was co-infiltrated with the nucleus marker NLS-mCherry in tobacco epidermal cells and observed under confocal microscopy. Bar = 50 μm.

**Table S1.**
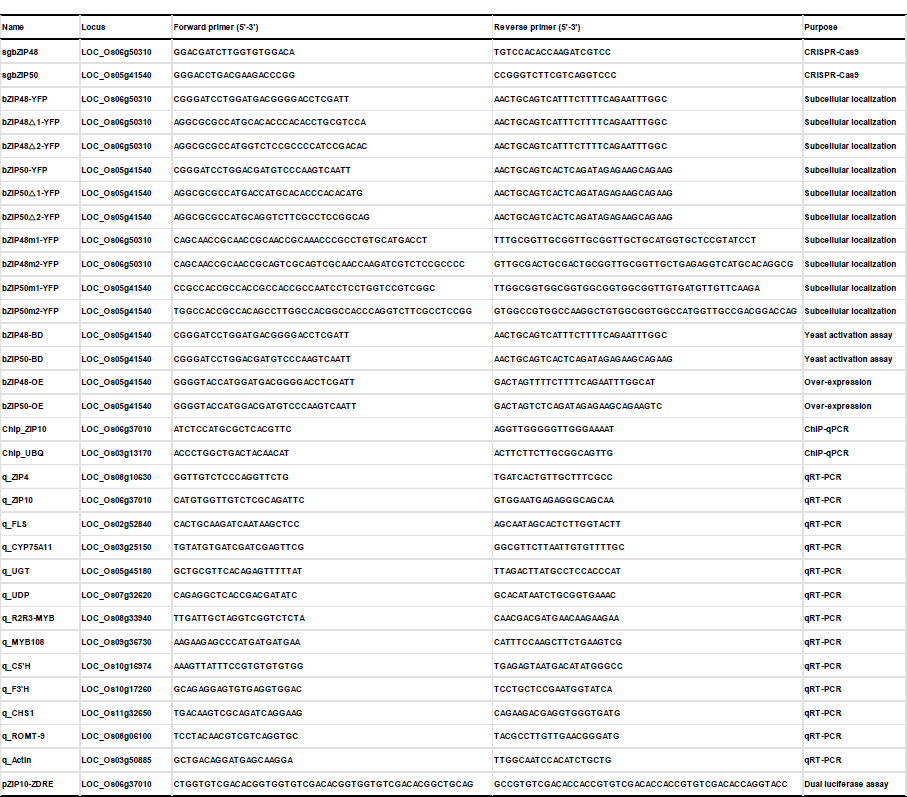
Primers Used in the Study.

Dataset S1. Differentially Expressed Genes under Zn Deficiency between Wild-type (ZH11) and *OsbZIP48*/*50* double mutant (*dm-1*).

## Notes

### Competing Interest Statement

The authors have declared no competing interest.

